# Imbalanced mitochondrial dynamics in human and mouse PD brains

**DOI:** 10.1101/2025.01.27.635175

**Authors:** Harry J. Brown, Rebecca Z. Fan, Riley Bell, Said S. Salehe, Carlos Martínez Martínez, Yanhao Lai, Kim Tieu

**Affiliations:** College of Arts and Sciences, Florida International University, Miami, FL; Department of Environmental Health Sciences, Florida International University, Miami, FL; Biomolecular Sciences Institute, Florida International University, Miami, FL; College of Biological Sciences, University of California, Davis, CA

**Keywords:** Parkinson’s disease, α-synuclein, dynamin-related protein 1, mitochondrial dynamics, protein aggregation, neurodegeneration

## Abstract

Mitochondrial dysfunction is a major pathogenic mechanism in Parkinson’s disease (PD). Emerging studies have shown that dysregulation in mitochondrial dynamics (fission/fusion/movement) has a major negative impact on mitochondria - both morphologically and functionally. Partial genetic deletion and pharmacological inhibition of the mitochondrial fission dynamin-related protein 1 (Drp1) have been demonstrated to be beneficial in experimental models of PD. However, the expression of DRP1 (and other fission and fusion genes/proteins) has not been investigated in the brains of Parkinson’s patients. Without these data, the question remains whether targeting DRP1 is a valid therapeutic target for PD. To address this gap of knowledge, first, we used post-mortem substantia nigra specimens of Parkinson’s patients and controls. Significant increases in the levels of both *DNM1L*, which encodes DRP1, as well as the DRP1 protein were detected in Parkinson’s patients. Immunostaining revealed increased DRP1 expression in dopamine (DA) neurons, astrocytes, and microglia. In addition to DRP1, the levels of other fission and fusion genes/proteins were also altered in Parkinson’s patients. To complement these human studies and given the significant role of α-synuclein in PD pathogenesis, we performed time-course studies (3-, 6- and 12-month) using transgenic mice overexpressing human wild-type *SNCA* under the mouse*Thy-1* promoter. As early as 6 months old, we detected an upregulation of *Dnm1l* and Drp1 in the nigral DA neurons of the *SNCA* mice as compared to their WT littermates. Furthermore, these mutant animals exhibited more Drp1 phosphorylation at serine 616, which promotes its translocation to mitochondria to induce fragmentation. Together, this study shows an upregulation of DRP1/Drp1 and alterations in other fission/fusion proteins in both human and mouse PD brains, leading to a pro-fission phenotype, providing additional evidence that blocking mitochondrial fission or promoting fusion is a potential therapeutic strategy for PD.

## Introduction

Parkinson’s disease (PD) is the second most prevalent and the fastest growing neurodegenerative disease [12]. This debilitating neurological disorder is characterized pathologically by the presence of toxic α-synuclein (α-syn) and loss of dopamine (DA) neurons in the substantia nigra compacta (SNpc). Since its first discovery almost three decades ago in familial PD [35], α-syn has increasingly received attention due to its prominent role, not only in familial but also in sporadic PD. The significance of this protein is best illustrated when it was recently reported to be a reliable biomarker for PD [37]. Aggregation of this small protein has long been found in the Lewy bodies (LBs) [38], which are intracellular inclusions considered to be the histological hallmark of PD. Although it is still a topic of debate whether LBs are neurotoxic or neuroprotective by sequestering the toxic oligomeric α-syn [5, 30], a recent study reported that widespread non-inclusion pathology of toxic α-syn oligomers occurs prior to the formation of LBs [18]. The neurotoxic effects of α-syn have also been demonstrated experimentally to spread from one cell to another in a prion-like fashion [31, 33]. Overall, abnormal conformation of α-syn is neurotoxic in PD.

Several pathogenic mechanisms associated with α-syn have been proposed, ranging from mitochondrial dysfunction, impaired protein degradation pathways (autophagy and ubiquitin proteasomal system), oxidative stress, synaptic dysfunction, and neuroinflammation. These mechanisms are not mutually exclusive and impairing one pathway such as mitochondrial function can result in activation of other pathogenic mechanisms [17]. The negative impact of α-syn has been demonstrated experimentally in both *in vitro* and *in vivo* models [3, 10, 11, 13, 40] by impairing mitochondrial respiration. These studies primarily focus on the impact of α-syn on the mitochondrial electron transport chain. However, some *in vivo* studies have also reported that α-syn overexpression induces mitochondrial fragmentation [3, 6], suggesting imbalance of mitochondrial dynamics leading to a pro-fission phenotype.

Mitochondrial dynamics refers to the continuous process of fission (division), fusion (joining) and movement of mitochondria. It is now recognized that a balance in mitochondrial fusion and fission is critical to neuronal function and viability. Fission and fusion are regulated by specific proteins. The outer mitochondrial mitofusin proteins (Mfn1 and Mfn2) and the inner mitochondrial protein optic atrophy 1 (OPA-1) are responsible for mitochondrial fusion. Mitochondrial fission is controlled by a separate set of proteins: Mitochondrial Fission Factor (Mff), Fission-1 (Fis1), as well as Mitochondrial Dynamics Proteins of 49 and 51 kDa (MiD49 and MiD51, respectively) are anchored to the OMM where they recruit cytosolic Dynamin-related protein-1 (Drp1), which then oligomerizes and forms a ring-like structure around the mitochondria to constrict and divide them [4, 21, 28]. Although the role of Drp1 in mitochondrial division is critical to normal cellular function, excessive mitochondrial fission is detrimental to cells. Accumulating evidence indicates that partial inhibition of Drp1 function is protective in experimental models of neurodegenerative diseases [1, 28]. However, the expression of Drp1 has not been investigated in the brain samples of Parkinson’s patients. This information is critical if this mitochondrial fission is to be considered as a potential therapeutic target for PD. In the present study we provide data showing the expression of DRP1 and other mitochondrial fission and fusion proteins/genes in the substantia nigra (SN) post-mortem samples of Parkinson’s patients. To complement these findings, we performed time-course studies in transgenic mice overexpressing human wild-type α-syn under the *Thy1* promoter [7, 34]. Together, these results show an upregulation of DRP1/Drp1 and alterations in other fission/fusion proteins leading to a pro-fission phenotype.

## Materials and Methods

### Animals

All animals used in this study were approved by the Institutional Animal Care and Use Committee at Florida International University. Mice expressing human *SNCA* under the *Thy-1* promoter [C57BL/6N-Tg(Thy1-SNCA)15Mjff/J, Strain #017682] are commercially available from the Jackson Laboratory. Hemizygotes were crossed with C57BL/6J (Jackson Laboratory, #000664) mice to establish and maintain the colony. Heterozygous Drp1-knockout mice were generated and maintained as previously described [14]. To generate double mutant mice with heterozygous overexpression of α-syn and heterozygous Drp1-knockout mice, we crossed *SNCA*^+/−^ mice with *Dnm1l*^+/−^ mice.

### Human Tissue

All postmortem samples used were obtained from the NIH NeuroBioBank. Inclusion criteria for individuals with PD are clinical diagnosis and neuropathology (Lewy bodies and dopaminergic neurodegeneration in the substantia nigra). Co-morbidities are excluded for both Parkinson’s patients and control subjects. For more information, please refer to Supplementary Table 1.

### Gel-Free Immunoblotting

Gel-free immunoblotting was performed with Jess™ system (ProteinSimple, Bio-Techne) using the 12-230 kDa Fluorescence Separation Module (cat. no. SM-FL004 ProteinSimple, Bio-Techne), as previously described [14]. Briefly, cytosolic fractions were isolated from frozen pulverized tissue by homogenization with cytoplasmic extraction buffer (10mM HEPES, 60mM KCl, 1mM EDTA, 0.075% IPEGAL, 1mM PMSF and 1mM DTT, pH 7.6). Incubation at 4°C, rotating for 10 mins prior to centrifugation at 1500 rpm to collect cytosolic supernatant. Micro BCA protein assay (23235, Thermo Fisher Scientific) was used to determine sample protein concentration, 1.5µg protein of human samples was loaded per capillary and 3µg of mouse samples. Primary antibodies used: Drp1 (Cell signaling, 8570S, 1:200), Drp1 616 (Cell signaling, 4494S, 1:25), MiD51 (Proteintech, 67808-1-Ig, 1:1000), Mfn2 (Sigma, M6319, 1:200), OPA-1 (BD transduction laboratories, 612606, 1:200).

### RNA extraction and Quantitative RT-PCR

Total RNA was extracted from human or mouse ventral midbrain using TRIzol (Thermo Fisher Scientific) per manufacturer’s instructions. RNA (500ng) was converted to complementary DNA (cDNA) using iScript Reverse Transcription Supermix (Bio-Rad). A QuantStudio 6 detection system (Thermo Fisher Scientific) was used for qPCR reaction using TaqMan assays and TaqMan Fast Advanced Master Mix (Thermo Fisher Scientific). Reaction conditions were 50°C for 2 minutes, 95°C for 2 minutes and 40 cycles of 95°C for 1s and 60°C for 20s. The 2^−ΔΔCT^ method was used for quantification. TaqMan assays used for human samples were *GAPDH* (HS02786624_g1), *DNM1L* (HS01552605_m1), *MFF* (HS00697394_g1), *MIEF2* (HS00541009_g1), *MIEF1* (HS01007730_g1), *OPA-1*: (HS01047013_m1), and *MFN2* (HS00208382_m1); for mouse studies were *Gapdh (*Mm99999914_g1), *Dnm1l* (Mm01342903_m1), *Mff* (Mm01273401_m1), *Mief1* (Mm00724569), *Mief2* (Mm01234249_g1), *Opa-1* (Mm01349707_g1), and *Mfn2* (Mm00500120_m1).

### Immunostaining

#### Human postmortem brain sections

Antigen retrieval: pre-mounted SN sections 10µm thick were deparaffinized overnight at 60°C. Rehydration was performed sequentially using: [100%] xylene, [100%] ethanol, [95%] ethanol, [70%] ethanol, [50%] ethanol and distilled water. Antigen retrieval was performed using Antigen Unmasking Solution (Vector, H-3300) and boiled in a steamer for 20 minutes before cooling at room temperature (RT) for 30 minutes.

Immunofluorescence: deparaffinized sections were washed in [1X] PBS, followed by permeabilization in [10%] Triton-X for 5 minutes. Sections were washed in [1X] PBS before incubation in [1%] Sudan black solution in [70%] ethanol for 2 hrs at RT. Subsequently sections were incubated overnight at 4 °C with primary antibody solution ([4%] normal goat serum (NGS), [0.5%] Triton-X, [0.05%] Tween-20, and [1X] PBS) contained within a humidity chamber. Sections were washed in [1X] PBS and incubated in secondary antibody solution for 1 hr at RT ([4%] NGS in [1X] PBS). Sections were washed in [1X] PBS and mounted using prolong gold antifade reagent with DAPI (Thermo Fisher Scientific, #P36941).

Immunohistochemistry: Brain sections were deparaffinized as described above and washed in [0.1M] TBS. Sections were then permeabilized and had endogenous peroxidases quenched with [0.3%] hydrogen peroxide in methanol for 10 minutes with shaking. Sections were washed in [0.1M] TBS before blocking for 1hr at RT using [5%] NGS in [0.1M] TBS. Tissue sections were then incubated in corresponding primary antibodies overnight at 4 °C in [0.1M] TBS containing [2%] NGS. Hereafter tissue sections were washed in [0.1M] TBS and incubated for 1hr at RT in corresponding biotinylated secondary antibody (1:200) in [0.1M] TBS with [2%] NGS. Sections were washed in 0.1M TBS before incubation for 1hr at RT in ABC solution according to manufacturer’s guidelines (Vector, PK6100). ABC solutions were made 30mins prior to usage. Sectionswere washed in [0.1M] TBS. Color generation was performed using DAB substrate kit, with nikel enhancement according to manufacturer’s guidelines (Vector, SK-4100). Sections were dehydrated in [95%] ethanol, [100%] ethanol and xylene before being sealed using DPX Mountant (Sigma, 06522). Primary antibodies used were DRP1 (BD biosciences, 6111133, 1:500), IBA1 (Wako, 019-19741, 1:1000), GFAP (Invitrogen, PA1-10004, 1:20,000).

#### Mouse brain sections

Mice were transcardially perfused, stored and sectioned as previously described [14]. Ventral midbrain sections (30 µm) were washed in [0.1M] TBS. Non-specific binding sites were blocked for 1.5hrs at RT using [4%] NGS and [0.5%] Triton-X in [0.1M] TBS. Primary antibodies were diluted in [2%] NGS, [0.3%] Triton-X in [0.1M] TBS and incubated overnight at 4°C whilst shaking. Sections were washed in [0.1M] TBS and incubated in secondary antibodies for 1h at RT whilst shaking. Primary antibodies: Drp1 (BD biosciences, 6111133, 1:500), Iba1 (Wako, 019-19741, 1:1000), GFAP (Invitrogen, PA1-10004, 1:5000), TH (Abcam, ab76442, 1:2000), and TOMM20 (Abcam, ab186735, 1:1000). Secondary antibodies: Alexa Fluro 488 goat anti-mouse IgG (H+L) (Invitrogen, A11029, 1:1000), Alexa Fluro 647 goat anti-chicken IgY (Invitrogen, 1016315-1, 1:1000), Alexa Fluro 568 goat anti-rabbit IgG (H+L) (Invitrogen, A11011, 1:1000).

### Mitochondrial Network Analysis

Confocal z-stacked images of TOMM20 immunostaining were used for mitochondrial network analysis as previously described [14]. Briefly, image analysis was performed in Fiji in this order: Z stacked images were merged into 2D format. Process, filters, unsharp mask; Process, enhance local contrast; Process, filters, median, binary, make binary; Process, binary, skeletonize; Analyze, skeleton, analyze 2D/3D skeleton. Hereafter the StuartLab plugin was used to analyze the mitochondrial network [41].

### IMARIS 3D rendering

Imaris (Oxford instruments) was used to determine Drp1 and mitochondrial colocalization. Imported confocal images had surfaces created and masked to visualize individual mitochondrial networks. The surfaces’ function was used to create a 3D render of tyrosine hydroxylase (TH)-positive neurons. Drp1 staining inside the 3D render of TH-positive neurons had the spots’ function applied to render Drp1 puncta. TOMM20 staining localized inside the 3D render of TH-positive neurons had the surfaces’ function applied to 3D render mitochondria. Individual mitochondria were separated using the split touching object’s function and labelled using individual object IDs. Proximity filtering of Drp1 spots touching mitochondrial renders yielded two Drp1 sub-populations: touching mitochondria or not touching mitochondria. Half of the diameter of Drp1 spots was used as a threshold to determine if Drp1 was touching or not touching mitochondria.

### Statistical Analysis

All data points are expressed as mean ± SEM and the mean differences between groups were compared using an unpaired t-test or the Two-Way Analysis of Variance (Two-Way ANOVA) as indicated in each figure. Data from the animal studies were analyzed using either an unpaired t-test for two-group comparison or a Two-Way ANOVA for mean difference comparison between two variables followed by Tukey’s post hoc testing. For human data, an unpaired t-test was used to compare the mean difference between Parkinson’s patients and controlsamples for different markers. To account for potential confounding effects of age and sex, regression analysis was implemented with age and sex as covariates using the *lm()* function of the stats package (for linear models) in the RStudio programming environment (R 4.4.2, RStudio 12.0 Build 467). Potential outlier data points were identified using a combination of Cook’s D, Standardized Residuals, and leverage from the regression results. Identified outlier data points were removed from the subsequent analysis.

## Results

### Drp1 is upregulated in the substantia nigra of Parkinson’s patients

Imbalanced mitochondrial dynamics have been reported in experimental models of PD [1, 22, 28]. However, a pro-fission/fusion phenotype in human Parkinson’s patients brain samples is yet to be established. To address this gap, we obtained thirty-two (16 with PD and 16 without PD) postmortem human SNpc samples from the NIH NeuroBioBank. Please refer to Supplementary Table 1 for more information about these human subjects. Given that partial Drp1 inhibition has been reported to be protective in experimental models of PD [3, 9, 13, 15, 24, 36, 39], we first addressed whether DRP1 was upregulated within the SNpc of postmortem samples of human Parkinson’s patients and whether there were any sex differences. The expression of DRP1 was assessed at both a transcriptional (mRNA) and translational (protein) level. As shown in Fig. 1a and 1b, significant increases in levels of both the gene that encodes DRP1, *Dynamin-1 like protein* (*DNM1L),* and DRP1 protein were seen in Parkinson’s patients compared with controls. No age and sex differences were detectable between control or Parkinson’s subjects (Supplementary Table 2) and therefore, data were pooled for analysis in Fig. 1a and 1b. To determine if increases in DRP1 were cell-type specific, we performed immunostaining for astrocytes, microglia and DA neurons (Fig. 1c-1e). DRP1 was expressed in all three cell types and consistent with the increased in gene and protein levels as seen in Fig. 1a and 1b, Parkinson’s patients had higher immunoreactivity of DRP1 in these cells. Together, these data indicate that DRP1 is significantly increased in Parkinson’s patients at both transcriptional and translational levels and is not apparent to be cell-type specific.

**Fig. 1.**
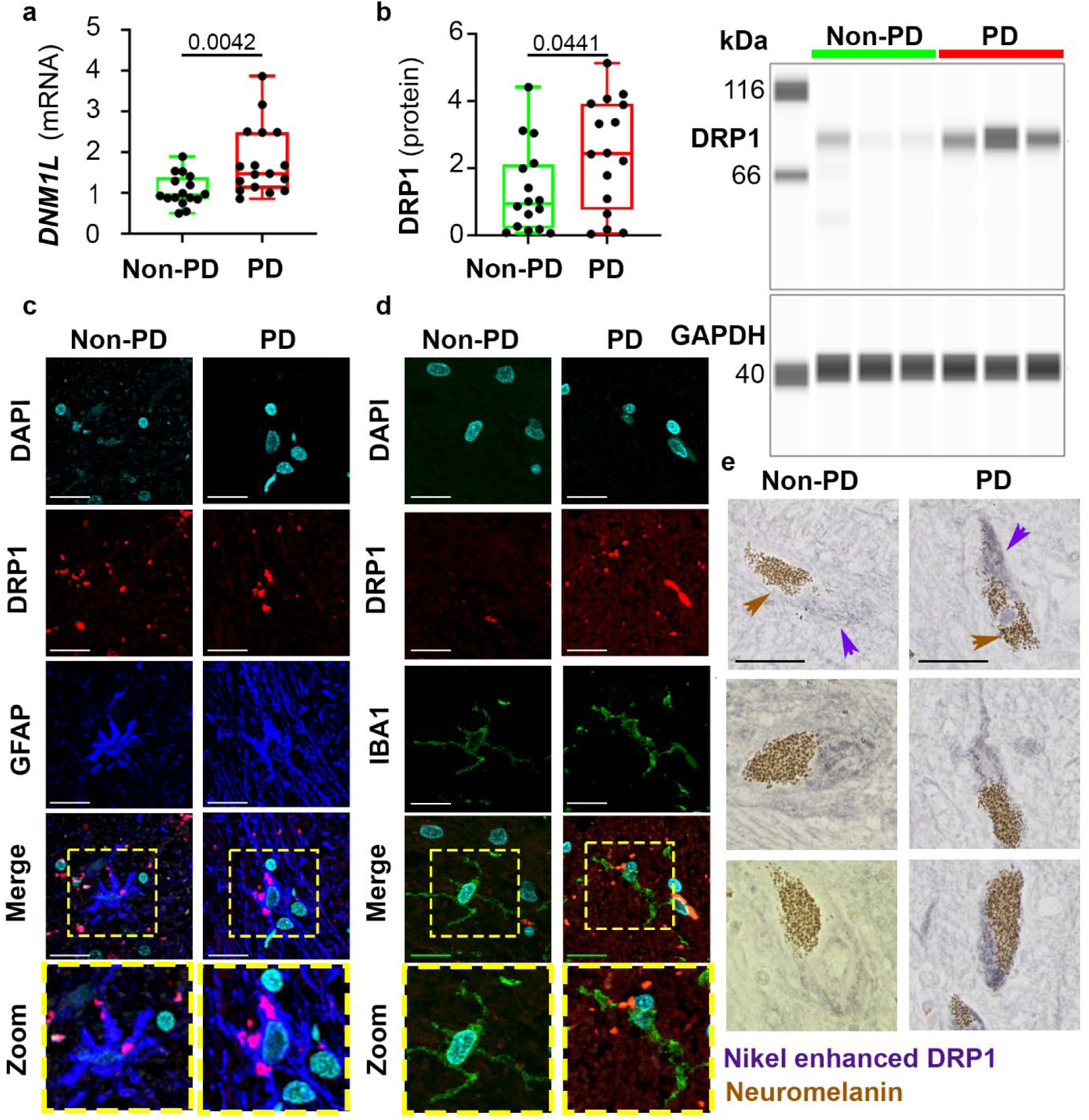
Increased DRP1 levels in human Parkinson’s patients brains. Post-mortem substantia nigra of PD and non-PD subjects were analyzed for the mRNA levels of *DNM1L* using qPCR **a**, and for the levels of DRP1 protein using gel-free immunoblotting **b**. Data were normalized to *GAPDH* mRNA and GAPDH protein, respectively. Immunofluorescence imaging of DRP1 levels in astrocytes **c**, and microglia **d.** To avoid autofluorescence from lipofuscin in DA neurons, immunohistochemistry using diaminobenzidine with nickel enhancement was performed to visualize DRP1 (purple arrows) in neuromelanin- (brown arrows) containing DA neurons **e.** Scale bar=20µm for **c-e**. Data represent mean ± SEM, unpaired t-test, n=16

### Alterations of other mitochondrial fission and fusion genes/proteins in the nigra of Parkinson’s patients

To have a more complete profile of whether or how other mitochondrial dynamics related genes and proteins were altered in Parkinson’s patients, we analyzed the expression of other mitochondrial fission and fusion mediators. Transcriptionally, *Mitochondrial Elongation Factor-1* (*MIEF1,* which encodes *MID51*) and *OPA-1* were significantly upregulated in Parkinsons’s patients (Fig. 2a) as compared to non-PD control subjects. *Mitochondrial Elongation Factor-2* (*MIEF2*), *Mitochondrial Fission Factor* (*MFF)* and *Mitofusin 2* (*MFN2*) levels were not significantly changed in Parkinson’s patients (Fig. 2a). Immunoblotting further confirmed the increases in Mitochondrial Dynamics Protein of 51kDa (MiD51) and OPA-1 protein levels in Parkinson’s patients (Fig. 2b). Overall, these data identify that both pro-fission and pro-fusion genes are differentially expressed in Parkinson’s patients. Such alterations in both fission and fusion factors likely result from a compensatory mechanism.

**Fig. 2.**
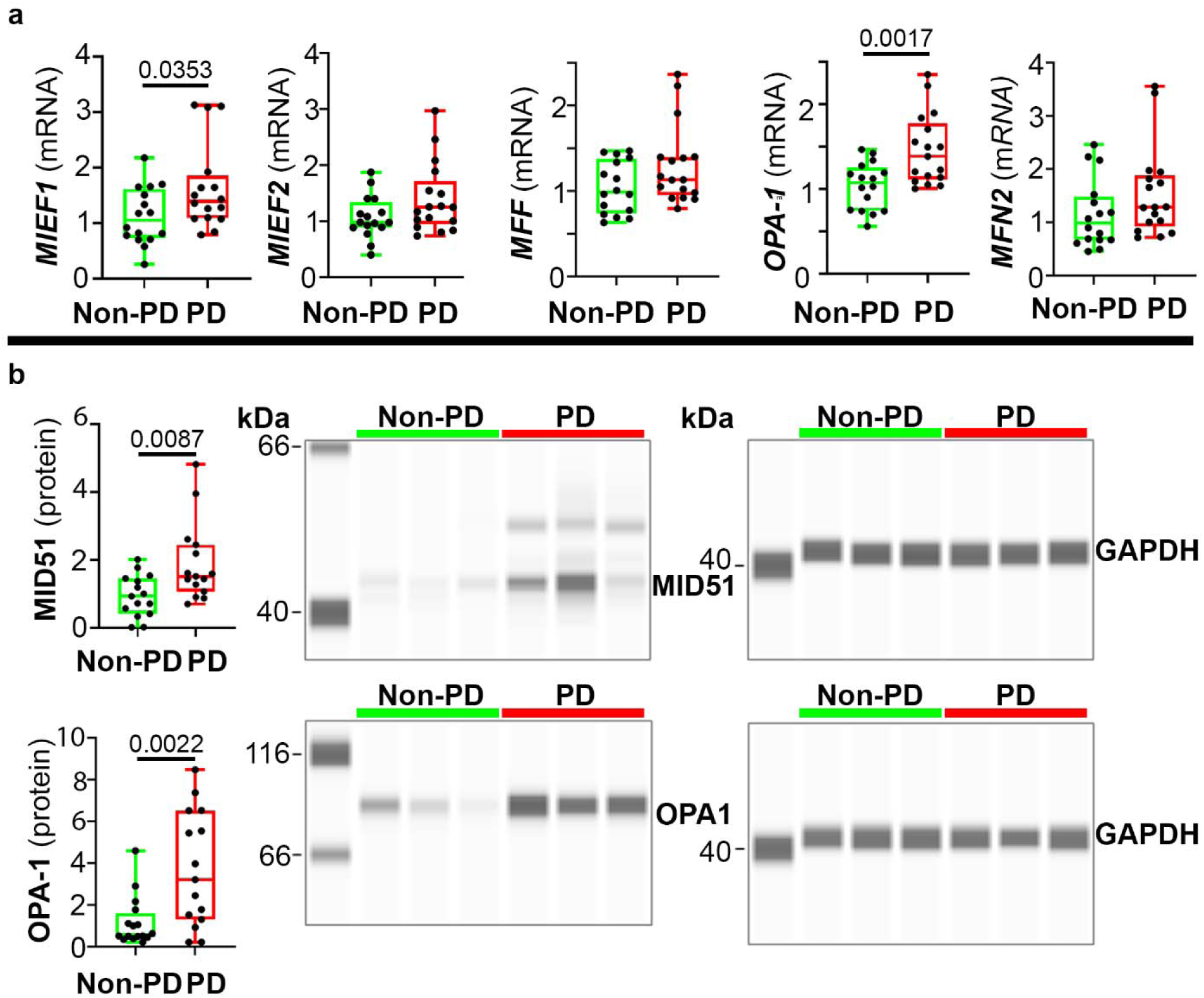
Levels of other mitochondrial dynamics genes and proteins in human brains. Post-mortem SN specimens of PD and non-PD subjects were analyzed for gene expression of *MIEF1*, *MIEF2*, *MFF, OPA-1*, and *MFN2* using qPCR **a**, or MID1 and OPA-1 proteins using gel-free immunoblotting **b**. Data represent mean ± SEM, normalized to *GAPDH* or GAPDH. Unpaired t-test, n=16

### Drp1 is transiently upregulated in the nigral DA neurons in a PD mouse model

Although translational, there are inherent limitations of using human post-mortem samples. Therefore, we turned to a transgenic mouse model of PD to further investigate the role of Drp1 in the presence of α-syn, a prominent protein involved in PD pathogenesis. With the C57BL/6N-Tg(Thy1-*SNCA*)15Mjff/J mice, which overexpress human *SNCA* under the *Thy1* promoter (*SNCA*^+/−^) [7, 34], we performed time-course studies at ages 3-, 6-, and 12-months. At 6-month-old, *Dnm1l* gene expression in the *SNCA* mice was upregulated when compared to age-matched WT littermate controls (Fig. 3a). We also observed that aging by itself increased *Dnml1* levels in the WT mice, but to a lesser extent than when combined with α-syn. To further investigate whether these changes also occurred in the nigral DA neurons, we performed immunofluorescence staining and detected higher immunoreactivity of Drp1 in TH-positive neurons in 6-month-old *SNCA* mice (Fig. 3b).

**Fig. 3.**
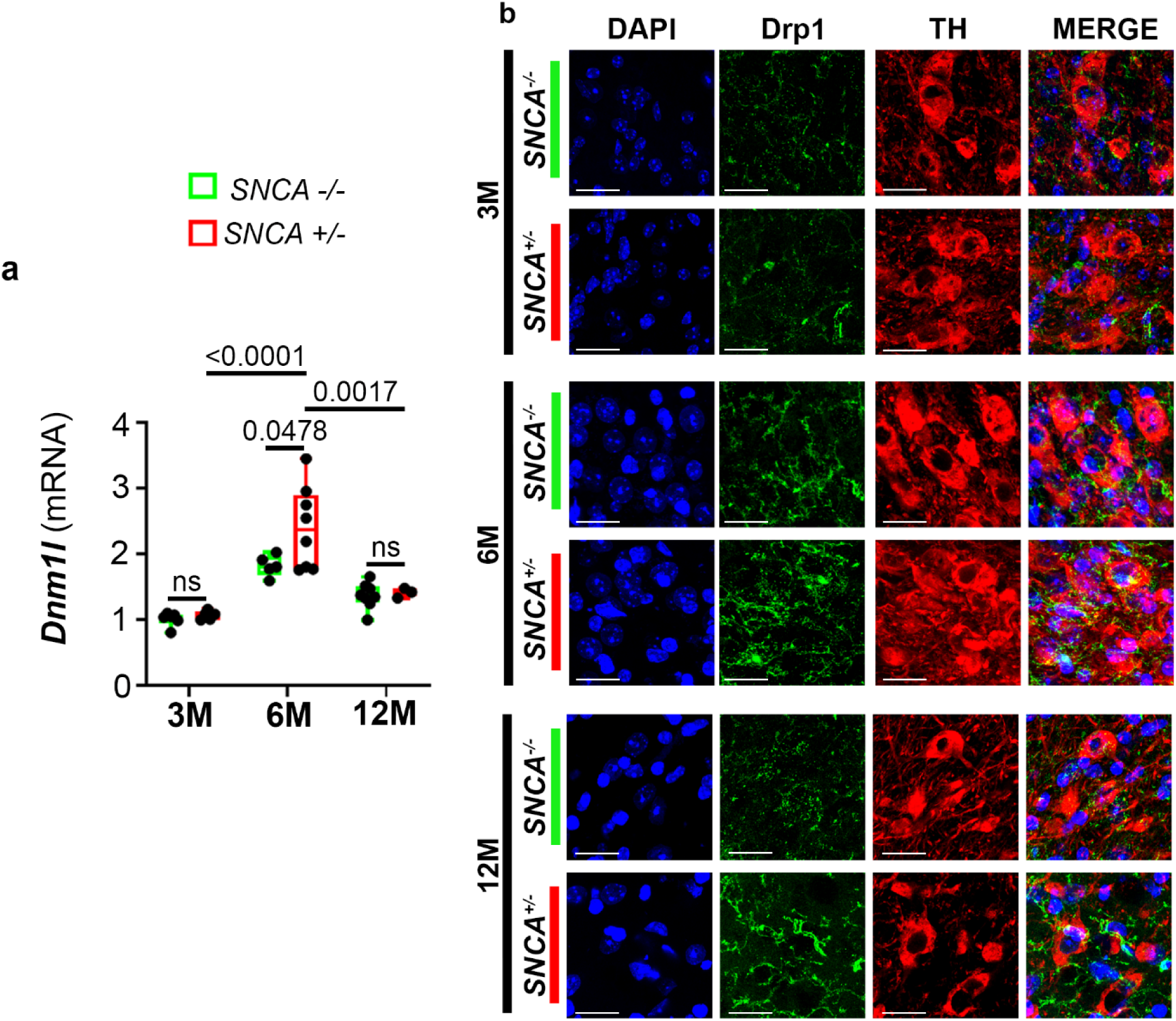
Longitudinal assessment of Drp1 levels in *SNCA* mice. **a** Ventral midbrains of heterozygous transgenic mice overexpressing human α-syn (*SNCA*^+/−^) and their WT littermates (*SNCA*^-/-^) at ages 3-, 6-, and 12-months were analyzed for mRNA levels of *Dnm1l* (normalized to *Gapdh*) using qPCR. **b** Immunofluorescence imaging of Drp1 colocalizing with TH-positive neurons of the SNpc. Scale bar=10µm. Data represent mean ± SEM. Age-dependent and genotype-dependent changes were analyzed using Two-Way ANOVA followed by Tukey’s post hoc tests. Genotypic changes independent of age were analyzed using an unpaired t-test, n=3-8

To corroborate changes in human Parkinson’s patients with the *SNCA* mouse model, we performed immunoblotting in 3-, 6-, and 12-month-old *SNCA* mice for other mitochondrial fission and fusion markers (MiD51, Mfn2, and Opa-1) as in the human samples (Supplementary Fig. S1). No significant genotypic changes were seen between WT and *SNCA* mice in these markers at any age. We did, however, observe age-dependent changes in MiD51 and Mfn2, both of them peaked in 6-month-old mice before decreasing in 12-month-old mice. Opa-1 levels significantly decreased with age. These findings highlight that overexpression of α-syn did not significantly alter the levels of MiD51, Mfn2, or Opa-1. Whereas age did significantly alter the levels of these proteins.

In combination, from this *in vivo* time-course study, we discovered that of all the fission and fusion markers that we assessed, Drp1 was most affected by α-syn, especially in the 6-month-old mice. The observations of normal levels of Drp1 at 12-months-old suggest that mice have a better ability to combat this increase as compared to Parkinson’s patients. However, this transient increase may be sufficient to induce a negative impact on mitochondria.

### *SNCA* mice exhibit age-dependent mitochondrial fragmentation

To further investigate whether increased Drp1 in the 6-month-old transgenic *SNCA* mice would result in a pro-fission phenotype, we co-labelled TH-positive neurons in the SNpc of these mice with translocase of outer mitochondrial membrane 20 (TOMM20) to visualize mitochondrial morphology. As shown in Fig. 4a, although mitochondrial fragmentation was observed in both genotypes in an age-dependent manner, *SNCA* mice exhibited more severe fragmentation, as evidenced by a decrease in branch length compared to their WT age-matched littermates (Fig. 4a). To confirm that these changes in mitochondrial morphology were in part attributed to Drp1, we replicated this experiment using 6-month-old double mutant mice with heterozygous overexpression of α-syn and heterozygous Drp1-knockout (Supplementary Fig. S2). As compared to littermates with only α-syn overexpression, double mutant mice exhibited longer mitochondrial branch length, indicative of a protection against the effects of α-syn. To obtain additional evidence that links Drp1 to mitochondrial fragmentation, we performed Drp1 colocalization analysis with mitochondria in TH-positive neurons of the SNpc in 6- and 12-month-old *SNCA* and WT littermates using the 3D rendering software IMARIS (Supplementary Fig. S3). This technique allowed us to clearly segregate Drp1 in contact with mitochondria from the mitochondrial absent Drp1 population. An increase in Drp1 translocating to mitochondria was detected in 6-month-old *SNCA* mice when compared to age-matched WT controls. Although statistically significant, the extent of Drp1 translocation in the 12-month-old *SNCA* mice is less(Fig. 4b).

**Fig. 4.**
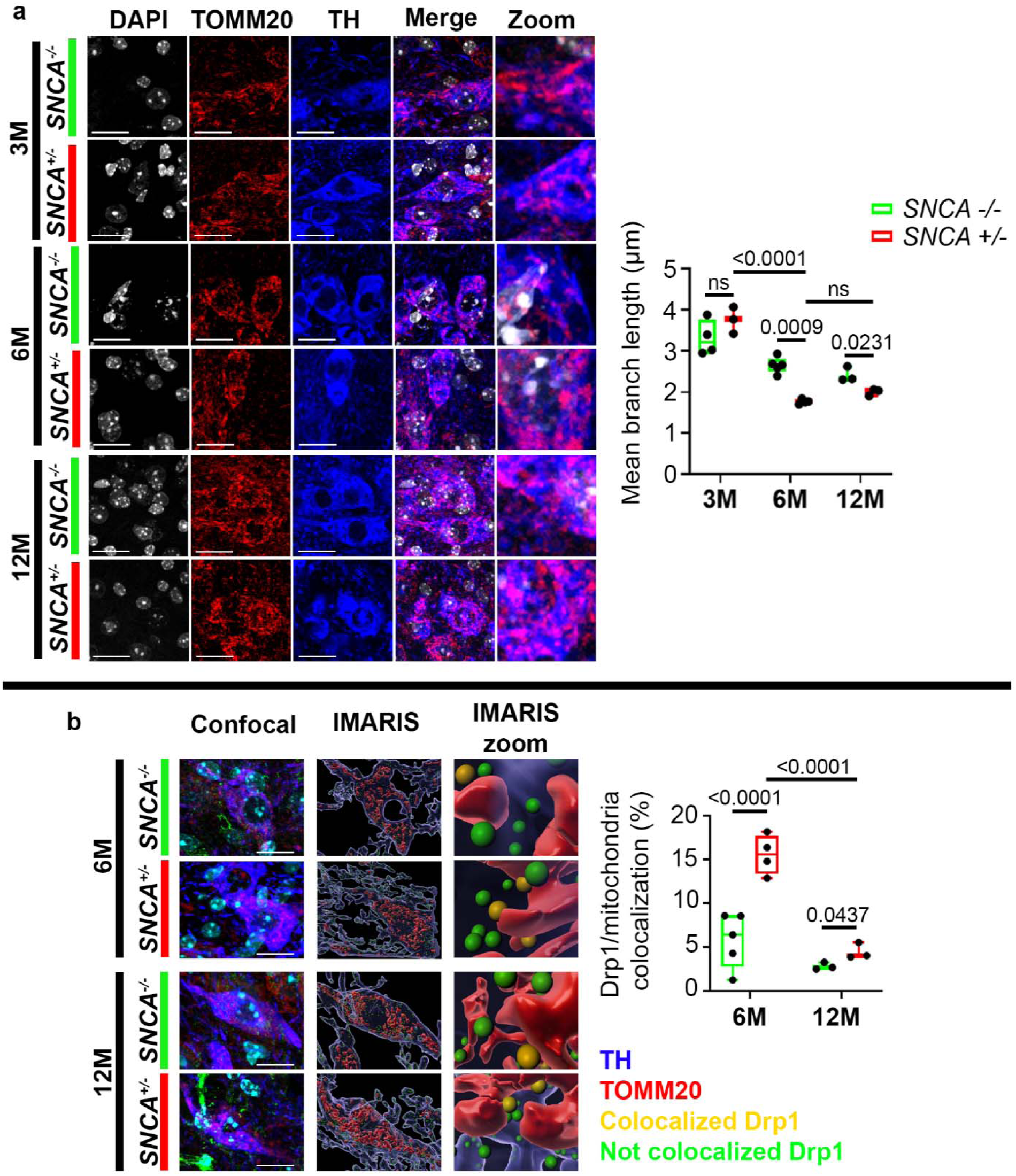
Age-dependent mitochondrial fragmentation in *SNCA* mice. **a** Immunofluorescence of TOMM20 to identify mitochondrial morphology in TH-positive neurons of the SNpc in 3-, 6-, and 12-month-old *SNCA* mice, scale bar=20µm. **b** Imaris 3D rendering of Drp1 mitochondrial colocalization in 6- and 12-month-old *SNCA* mice. Individual immunofluorescent channels from confocal images were outlined to isolate TH-positive neurons. Zoomed images of IMARIS display IMARIS 3D renders of Drp1 colocalization on mitochondria, Scale bar=10 µm. Data represent mean ± SEM. Age dependent and genotype dependent changes were analyzed using Two-Way ANOVA followed by Tukey’s post hoc tests. Genotypic changes independent of age were analyzed using unpaired t-test, n=3-5

Overall, these data demonstrate that there is a significant increase in the expression of Drp1 and its translocation to mitochondria, resulting in severe fragmentation in the nigral TH-positive neurons of the *SNCA* mice, starting at 6-month-old. Additionally, the effects of aging are further exacerbated by the presence of α-syn.

### Post-translational modification of Drp1 is increased in *SNCA* mice

One well-established post-translational modification that promotes Drp1 translocation to mitochondria to induce fragmentation is its phosphorylation at the serine 616 residue (pDrp1-S616) [4, 27]. To investigate whether pDrp1-s616 played a role in mitochondrial fragmentation in the *SNCA* mice, we performed immunofluorescence for pDrp-s616 in the nigral TH-positive neurons, followed by quantification using fluorescent intensity (Fig. 5). No differences in pDrp-s616 were detectable between 3-month-old *SNCA* and WT mice. However, pDrp1-s616 was significantly increased in TH-positive neurons of 6-month-old *SNCA* mice, when compared to WT age-matched controls. Although not as extensive, increased pDrp1-s616 was also observed in 12-month-old *SNCA* mice. In aggregate, these results indicate that α-syn promotes post-translational phosphorylation of Drp1 at a site that promotes its translocation to mitochondria to induce mitochondrial fragmentation.

**Fig. 5.**
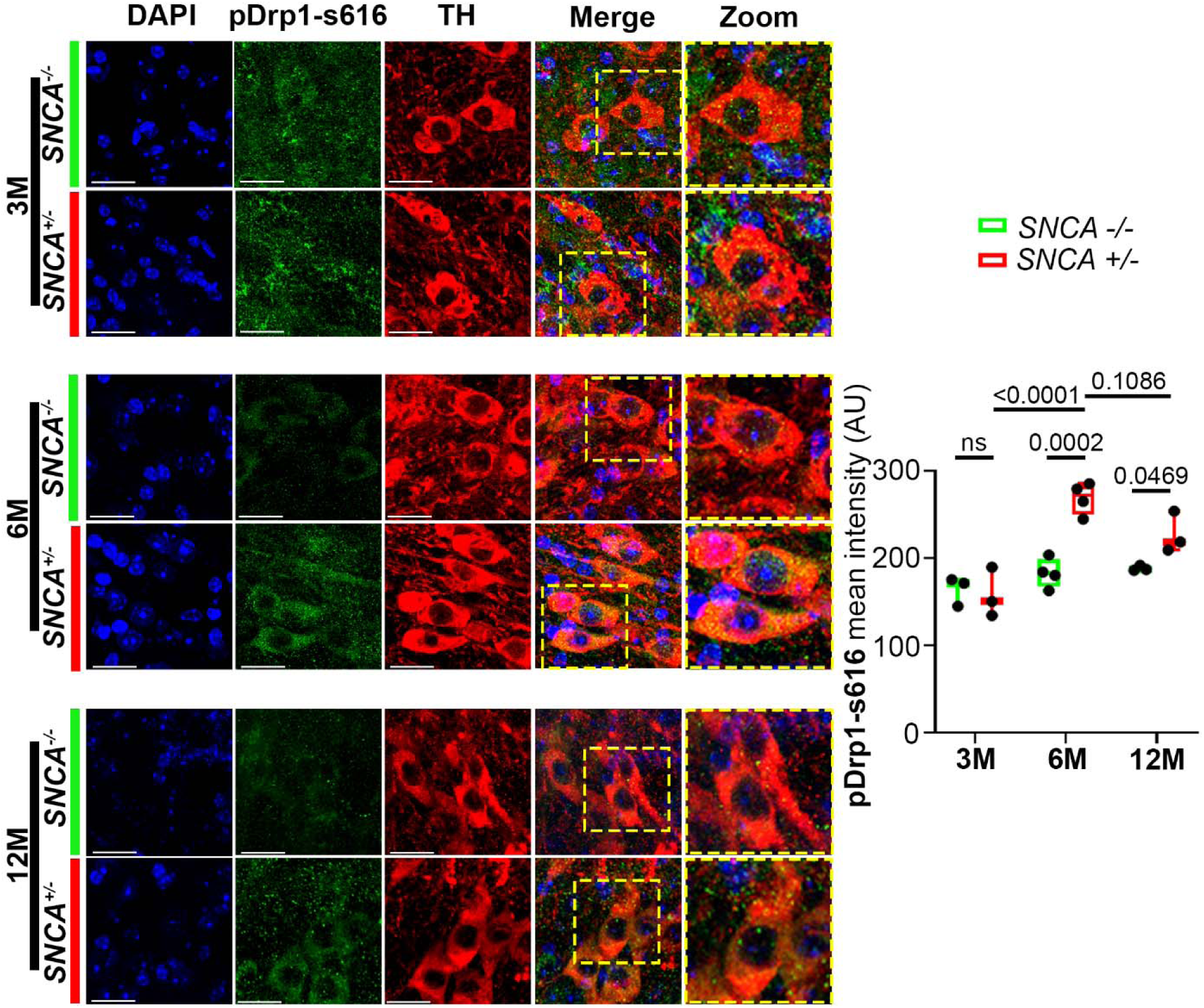
pDrp1-s616 is enriched in TH positive neurons of 6- and 12-month-old *SNCA* mice. Immunofluorescence of pDrp1-s616 in TH-positive neurons of the SNpc. Fluorescent intensity was quantified (AU: arbitrary unit). Data represent mean ± SEM. Age dependent and genotype dependent changes were analyzed using Two-Way ANOVA followed by Tukey’s post hoc tests. Genotypic changes independent of age were analyzed using unpaired t-test, n=3-5

## Discussion

Balanced mitochondrial fission and fusion is essential to cellular function and viability. Drp1 plays a pivotal role in maintaining this balance by mediating fission, a process critical to the removal of damaged mitochondria through mitophagy and the redistribution of mitochondrial content to meet localized cellular energy demands. Aberrant Drp1 activity, whether excessive or insufficient, has been reported to induce abnormal mitochondrial morphology, mitochondrial distribution, bioenergetic deficits, oxidative stress, apoptosis, and impaired autophagy-all of which may contribute to neurodegeneration and other diseases [4, 22, 27, 28]. Excessive mitochondrial fission with strategies used to reduce Drp1 function has been reported to be protective in experimental models of neurodegenerative diseases such as Alzheimer’s disease (AD) and PD [28]. In preclinical models of PD, pharmacological inhibitors (such as P110 and mdivi-1) and genetic manipulations (loss of Drp1 function or overexpression of fusion proteins) attenuate mitochondrial abnormalities and improve neuronal function and survival [3, 9, 13, 15, 24, 36, 39]. Together, such studies indicate DRP1 as a potential therapeutic target for PD. However, to date, the expression of this protein and other mitochondrial fission/fusion proteins have not been examined in the brains of Parkinson’s patients.

In the present study, we analyzed human post-mortem SNpc from 16 PD and 16 non-PD subjects for the expression of DRP1. Our data indicate that DRP1 is significantly increased in human Parkinson’s patients at both mRNA and protein levels. Furthermore, in these patients, immunoreactivity of DRP1 was higher in DA neurons, microglia and astrocytes, compared to those in non-PD individuals. Considering other studies have reported that blocking Drp1 reduces neuroinflammation [19, 29], our observations suggest reducing DRP1 levels may also attenuate neuroinflammation in PD, providing an additional neuroprotective mechanism. In addition to DRP1, we assessed the levels of other fission (*MIEF1*, *MIEF2*, *MFF*) and fusion (*OPA-1* and *MFN2*) genes. Consistent with the increased expression of *MIEF1* and *OPA1* transcripts, we also detected an increase in their corresponding gene products MiD51 and OPA1. Because MiD51 is a receptor for DRP1 [32], their elevated levels would result in pro-fission. Perhaps OPA1 is upregulated to compensate for this effect. Interestingly, we did not detect a similar change with another fusion gene *MFN2*, which has been reported to be reduced in patients with AD [43]. Given the variance in age and sex between Parkinson’s patients and non-PD controls, samples were adjusted for age and sex. No sex differences were identified between any of the analyzed mitochondrial fission/fusion proteins. Age also did not significantly influence the levels of these proteins. However, this undetectable age-dependent effect may be due to the lack of the longitudinal aging aspect in our patient population, whose ages ranged only from 75-88 years old. A younger cohort would be required to evaluate if the alterations in mitochondrial fission/fusion proteins are age dependent. Taken together, both pro-fission (DRP1 and MiD51) and pro-fusion (OPA-1) proteins were upregulated in Parkinson’s patients. One approach to determine whether such alterations would result in pro-fission or fusion phenotype is to quantify mitochondrial morphology. However, assessing mitochondrial morphology in human post-mortem samples is inherently challenging due to the rapid structural alterations that occur after death when organelles are assumed to cease functionality [2]. Given that mitochondria are highly dynamic organelles, their morphology can be influenced by factors such as agonal state, post-mortem interval, and the tissue preservation methods, making it difficult to distinguish pathological changes to the mitochondria from artifacts [2, 16].

To complement some of the inherent challenges such as the ones discussed above when using human brain samples and other concerns such as potential alterations to tissue integrity due to variability in post-mortem interval, heterogeneity of the severity of the disease, sporadic versus genetic PD, and potential confounding factors such as medications taken and exposure to environmental toxins, we included a transgenic mouse model overexpressing human WT α-syn, which is involved in both familial and sporadic PD. Another advantage of using the *SNCA* mice is that these animals allowed us the feasibility to directly study the *in vivo* effects of α-syn on mitochondrial dynamics in an age- and sex-dependent manner. Our results show an upregulation of Drp1 in the 6-month-old *SNCA* mice when compared to other age groups and their age-matched WT littermate controls. Interestingly, we observed that aging by itself also increased the levels of this fission protein, but to a lesser extent than when combined with α-syn. It is tempting to speculate that perhaps such alteration contributes to the age-dependent onset of PD. Mitochondrial fragmentation was observed in 6- and 12-month-old *SNCA* mice which was corroborated by increased mitochondrial colocalization of Drp1. These changes were observed in tandem with increases of pDrp1-s616 in TH-positive neurons of the SNpc in 6- and 12-month-old *SNCA* mice. The contribution of Drp1 to this phenotype was further corroborated whereby mitochondrial fragmentation was prevented in the Drp1-KO mice (Supplementary Fig. 2). We also observed dynamic changes in other pro-fission and fusion genes/proteins in the *SNCA* mice. Increased expression of pro-fission proteins, such as Drp1 and MiD51, may lead to a compensatory change in the regulation of pro-fusion proteins such as Mfn1, Mfn2, and Opa-1 to maintain mitochondrial homeostasis and prevent pathological fragmentation. Alterations in both fission and fusion proteins have also been reported in AD patients and experimental models of AD [20, 26, 43].

No changes were detectable in any mitochondrial proteins between 3-month-old WT and *SNCA* mice. It is possible that at this early age α-syn pathology is still developing. Although total α-syn levels in these *SNCA* mice increase at 3-months-old, a minor increase in α-syn pS129 levels only began in 8- and 12-month-old mice [7, 34]. It has been shown that in human Parkinson’s patients pathogenic aggregates of α-syn pS129 preferentially bind to mitochondria in contrast to their soluble counterparts [23, 42]. Specifically, the α-syn N-terminus binds to lipids on the mitochondrial membrane, such as cardiolipin [25]. Under pathophysiological conditions cardiolipin oxidizes and translocates from the inner to outer mitochondrial membrane [8]. Certain post-translationally modified species of α-syn bind with high affinity to the TOMM20, resulting in preventing the interaction of this mitochondrial receptor with its co-receptor, TOMM22, and impaired mitochondrial protein import [11]. Thus, these interactions with α-syn can impact the import and export of proteins and ions at the mitochondria leading to mitochondrial stress and potentially fragmentation. Such previous studies combined with the data we show of increased pDrp1-s616 phosphorylation and subsequent mitochondrial fragmentation in PD models highlight the important role that α-syn may play in mediating Drp1-induced mitochondrial fragmentation in the older *SNCA* mice.

## Conclusion

To our knowledge this is the first study that investigates mitochondrial fission and fusion factors in human Parkinson’s patients postmortem specimens, coupled with a transgenic mouse model overexpressing human α-syn. Our results highlight a dysregulation of mitochondrial fission and fusion proteins in Parkinson’s patients and *SNCA* mice. Furthermore, increased DRP1/Drp1 levels were consistently observed in these patients and mutant mice. Relevant to this study, as discussed, blocking Drp1 has been reported to be neuroprotective against mitochondrial dysfunction, oxidative stress, impaired autophagy, aggregated α-syn, and synaptic dysfunction in PD models. The present study highlights DRP1 as an attractive target for PD and synucleinopathies.

## Declarations

### Author contributions

KT and HJB contributed to the study conception and design. Material preparation was performed by HJB, RZF, SSS, and YL. Data collection and analysis were performed by HJB, RB, SSS, and CMM. HJB and KT were involved in drafting and revision of the manuscript. All other authors commented on previous versions of the manuscript and approved the final version.

### Ethics approval

#### Animal study

All mice in this study were bred, maintained, and characterized at animal care facility of Florida International University (FIU). Animal care and procedures were approved and conducted in accordance with the Institutional Animal Care and Use Committee at FIU (No. IACUC-24-088)

#### Human samples

This study utilized human tissue that was procured via NIH NeuroBioBank, which provides de-identified samples. This research was reviewed and deemed exempt by our Florida International University Institutional Review Board. The NIH NeuroBank protocols are in accordance with the ethical standards of our institution and with the 1964 Helsinki declaration and its later amendments or comparable ethical standards.

### Consent for publication

All authors have read the manuscript and accepted responsibility for the manuscript’s content.

### Materials and data availability

Human post-mortem samples are available through NIH-NeuroBiobank. Transgenic mice expressing human *SNCA* under the *Thy-1* promoter [C57BL/6N-Tg(Thy1-SNCA)15Mjff/J, Strain #017682] are commercially available from the Jackson Laboratory.

Data is provided within the manuscript or supplementary information files

## References

1 Archer SL (2013) Mitochondrial dynamics--mitochondrial fission and fusion in human diseases. N Engl J Med 369: 2236–2251

2 Barksdale KA, Perez-Costas E, Gandy JC, Melendez-Ferro M, Roberts RC, Bijur GN (2010) Mitochondrial viability in mouse and human postmortem brain. FASEB J 24: 3590–3599 Doi 10.1096/fj.09-152108

3 Bido S, Soria FN, Fan RZ, Bezard E, Tieu K (2017) Mitochondrial division inhibitor-1 is neuroprotective in the A53T-alpha-synuclein rat model of Parkinson’s disease. Sci Rep 7: 7495 Doi 10.1038/s41598-017-07181-0

4 Chan DC (2020) Mitochondrial Dynamics and Its Involvement in Disease. Annu Rev Pathol 15: 235–259 Doi 10.1146/annurev-pathmechdis-012419-032711

5 Chartier S, Duyckaerts C (2018) Is Lewy pathology in the human nervous system chiefly an indicator of neuronal protection or of toxicity? Cell Tissue Res 373: 149–160 Doi 10.1007/s00441-018-2854-6

6 Chen L, Xie Z, Turkson S, Zhuang X (2015) A53T human alpha-synuclein overexpression in transgenic mice induces pervasive mitochondria macroautophagy defects preceding dopamine neuron degeneration. Journal of Neuroscience, City, pp 890–905

7 Choi I, Zhang Y, Seegobin SP, Pruvost M, Wang Q, Purtell K, Zhang B, Yue Z (2020) Microglia clear neuron-released alpha-synuclein via selective autophagy and prevent neurodegeneration. Nat Commun 11: 1386 Doi 10.1038/s41467-020-15119-w

8 Chu CT, Ji J, Dagda RK, Jiang JF, Tyurina YY, Kapralov AA, Tyurin VA, Yanamala N, Shrivastava IH, Mohammadyani Det al (2013) Cardiolipin externalization to the outer mitochondrial membrane acts as an elimination signal for mitophagy in neuronal cells. Nat Cell Biol 15: 1197–1205 Doi 10.1038/ncb2837

9 Cui M, Tang X, Christian WV, Yoon Y, Tieu K (2010) Perturbations in mitochondrial dynamics induced by human mutant PINK1 can be rescued by the mitochondrial division inhibitor mdivi-1. J Biol Chem 285: 11740–11752

10 Devi L, Raghavendran V, Prabhu BM, Avadhani NG, Anandatheerthavarada HK (2008) Mitochondrial import and accumulation of alpha-synuclein impair complex I in human dopaminergic neuronal cultures and Parkinson disease brain. J Biol Chem 283: 9089–9100

11 Di-Maio R, Barrett PJ, Hoffman EK, Barrett CW, Zharikov A, Borah A, Hu X, McCoy J, Chu CT, Burton EA, et al (2016) alpha-Synuclein binds to TOM20 and inhibits mitochondrial protein import in Parkinson’s disease. Sci Transl Med 8: 342ra378

12 Dorsey ER, Sherer T, Okun MS, Bloem BR (2018) The Emerging Evidence of the Parkinson Pandemic. J Parkinsons Dis 8: S3–s8 Doi 10.3233/jpd-181474

13 Fan RZ, Guo M, Luo S, Cui M, Tieu K (2019) Exosome release and neuropathology induced by alpha-synuclein: new insights into protective mechanisms of Drp1 inhibition. Acta neuropathologica communications 7: 184 Doi 10.1186/s40478-019-0821-4

14 Fan RZ, Sportelli C, Lai Y, Salehe SS, Pinnell JR, Brown HJ, Richardson JR, Luo S, Tieu K (2024) A partial Drp1 knockout improves autophagy flux independent of mitochondrial function. Mol Neurodegener 19: 26 Doi 10.1186/s13024-024-00708-w

15 Filichia E, Hoffer B, Qi X, Luo Y (2016) Inhibition of Drp1 mitochondrial translocation provides neural protection in dopaminergic system in a Parkinson’s disease model induced by MPTP. Sci Rep, City, pp 32656

16 Harish G, Venkateshappa C, Mahadevan A, Pruthi N, Bharath MM, Shankar SK (2013) Mitochondrial function in human brains is affected by pre- and post mortem factors. Neuropathol Appl Neurobiol 39: 298–315 Doi 10.1111/j.1365-2990.2012.01285.x

17 Helley MP, Pinnell J, Sportelli C, Tieu K (2017) Mitochondria: A Common Target for Genetic Mutations and Environmental Toxicants in Parkinson’s Disease. Front Genet 8: 177 Doi 10.3389/fgene.2017.00177

18 Jensen NM, Fu Y, Betzer C, Li H, Elfarrash S, Shaib AH, Krah D, Vitic Z, Reimer L, Gram Het al (2024) MJF-14 proximity ligation assay detects early non-inclusion alpha-synuclein pathology with enhanced specificity and sensitivity. NPJ Parkinsons Dis 10: 227 Doi 10.1038/s41531-024-00841-9

19 Joshi AU, Minhas PS, Liddelow SA, Haileselassie B, Andreasson KI, Dorn GW, 2nd, Mochly-Rosen D (2019) Fragmented mitochondria released from microglia trigger A1 astrocytic response and propagate inflammatory neurodegeneration. Nat Neurosci 22: 1635–1648 Doi 10.1038/s41593-019-0486-0

20 Kandimalla R, Manczak M, Pradeepkiran JA, Morton H, Reddy PH (2021) A partial reduction of Drp1 improves cognitive behavior and enhances mitophagy, autophagy and dendritic spines in a transgenic tau mouse model of Alzheimer disease. Hum Mol Genet: Doi 10.1093/hmg/ddab360

21 Knott AB, Bossy-Wetzel E (2008) Impairing the mitochondrial fission and fusion balance: a new mechanism of neurodegeneration. 2008/12/17: 283–292

22 Knott AB, Perkins G, Schwarzenbacher R, Bossy-Wetzel E (2008) Mitochondrial fragmentation in neurodegeneration. Nat Rev Neurosci 9: 505–518

23 Ludtmann MHR, Angelova PR, Horrocks MH, Choi ML, Rodrigues M, Baev AY, Berezhnov AV, Yao Z, Little D, Banushi Bet al (2018) alpha-synuclein oligomers interact with ATP synthase and open the permeability transition pore in Parkinson’s disease. Nat Commun 9: 2293 Doi 10.1038/s41467-018-04422-2

24 Lutz AK, Exner N, Fett ME, Schlehe JS, Kloos K, Laemmermann K, Brunner B, Kurz-Drexler A, Vogel F, Reichert AS et al (2009) Loss of parkin or PINK1 function increases DRP1-dependent mitochondrial fragmentation. J Biol Chem 283: 22938–22951

25 Maldonado Vidaurri E, Chavez-Montes A, Garza Tapia M, Castro-Rios R, Gonzalez-Horta A (2018) Differential interaction of alpha-synuclein N-terminal segment with mitochondrial model membranes. Int J Biol Macromol 119: 1286–1293 Doi 10.1016/j.ijbiomac.2018.08.049

26 Manczak M, Kandimalla R, Fry D, Sesaki H, Reddy PH (2016) Protective effects of reduced dynamin-related protein 1 against amyloid beta-induced mitochondrial dysfunction and synaptic damage in Alzheimer’s disease. Hum Mol Genet: ddw330

27 Oettinghaus B, Licci M, Scorrano L, Frank S (2012) Less than perfect divorces: dysregulated mitochondrial fission and neurodegeneration. Acta Neuropathol 123: 189–203 Doi 10.1007/s00401-011-0930-z

28 Oliver D, Reddy PH (2019) Dynamics of Dynamin-Related Protein 1 in Alzheimer’s Disease and Other Neurodegenerative Diseases. Cells 8: Doi 10.3390/cells8090961

29 Park J, Choi H, Min JS, Park SJ, Kim JH, Park HJ, Kim B, Chae JI, Yim M, Lee DS (2013) Mitochondrial dynamics modulate the expression of pro-inflammatory mediators in microglial cells. J Neurochem 127: 221–232 Doi 10.1111/jnc.12361

30 Parkkinen L, O’Sullivan SS, Collins C, Petrie A, Holton JL, Revesz T, Lees AJ (2011) Disentangling the relationship between lewy bodies and nigral neuronal loss in Parkinson’s disease. J Parkinsons Dis 1: 277–286 Doi 10.3233/JPD-2011-11046

31 Peng C, Trojanowski JQ, Lee VM (2020) Protein transmission in neurodegenerative disease. Nat Rev Neurol 16: 199–212 Doi 10.1038/s41582-020-0333-7

32 Pernas L, Scorrano L (2016) Mito-Morphosis: Mitochondrial Fusion, Fission, and Cristae Remodeling as Key Mediators of Cellular Function. Annual review of physiology 78: 505–531 Doi 10.1146/annurev-physiol-021115-105011

33 Pinnell JR, Cui M, Tieu K (2021) Exosomes in Parkinson disease. J Neurochem 157: 413–428 Doi 10.1111/jnc.15288

34 Polinski NK, Martinez TN, Ramboz S, Sasner M, Herberth M, Switzer R, Ahmad SO, Pelligrino LJ, Clark SW, Marcus JN et al (2022) The GBA1 D409V mutation exacerbates synuclein pathology to differing extents in two alpha-synuclein models. Dis Model Mech 15: Doi 10.1242/dmm.049192

35 Polymeropoulos MH, Lavedan C, Leroy E, Ide SE, Dehejia A, Dutra A, Pike B, Root H, Rubenstein J, Boyer Ret al (1997) Mutation in the α-Synuclein Gene Identified in Families with Parkinson’s Disease. Science 276: 2045–2047 Doi doi:10.1126/science.276.5321.2045

36 Rappold PM, Cui M, Grima JC, Fan RZ, de Mesy-Bentley KL, Chen L, Zhuang X, Bowers WJ, Tieu K (2014) Drp1 inhibition attenuates neurotoxicity and dopamine release deficits in vivo. Nat Commun 5:5244. doi: 10.1038/ncomms6244:

37 Siderowf A, Concha-Marambio L, Lafontant DE, Farris CM, Ma Y, Urenia PA, Nguyen H, Alcalay RN, Chahine LM, Foroud Tet al (2023) Assessment of heterogeneity among participants in the Parkinson’s Progression Markers Initiative cohort using alpha-synuclein seed amplification: a cross-sectional study. Lancet Neurol 22: 407–417 Doi 10.1016/S1474-4422(23)00109-6

38 Spillantini MG, Schmidt ML, Lee VMY, Trojanowski JQ, Jakes R, Goedert M (1997) α-Synuclein in Lewy bodies. Nature 388: 839–840 Doi 10.1038/42166

39 Su YC, Qi X (2013) Inhibition of excessive mitochondrial fission reduced aberrant autophagy and neuronal damage caused by LRRK2 G2019S mutation. Hum Mol Genet 22: 4545–4561

40 Subramaniam SR, Vergnes L, Franich NR, Reue K, Chesselet MF (2014) Region specific mitochondrial impairment in mice with widespread overexpression of alpha-synuclein. Neurobiol Dis 70: 204–213

41 Valente AJ, Maddalena LA, Robb EL, Moradi F, Stuart JA (2017) A simple ImageJ macro tool for analyzing mitochondrial network morphology in mammalian cell culture. Acta Histochem 119: 315–326 Doi 10.1016/j.acthis.2017.03.001

42 Wang X, Becker K, Levine N, Zhang M, Lieberman AP, Moore DJ, Ma J (2019) Pathogenic alpha-synuclein aggregates preferentially bind to mitochondria and affect cellular respiration. Acta neuropathologica communications 7: 41 Doi 10.1186/s40478-019-0696-4

43 Wang X, Su B, Lee HG, Li X, Perry G, Smith MA, Zhu X (2009) Impaired balance of mitochondrial fission and fusion in Alzheimer’s disease. J Neurosci 29: 9090–9103

