## Supplementary information for "Imbalanced mitochondrial dynamics in human and mouse PD brains"

**Table 1**. Human postmortem SNpc sample information.

| **Sex** | **Age** | **Race** | **Postmortem interval (hrs)** | **Clinical/pathological Diagnosis** |
| --- | --- | --- | --- | --- |
| Female | 85 | White | 5 | Non-PD |
| Female | 75 | Black or African | 6.5 | Non-PD |
| Female | 76 | Unknown | 4.25 | Non-PD |
| Female | 86 | White | 4.38 | Non-PD |
| Female | 79 | White | 3.02 | Non-PD |
| Female | 85 | White | 5.27 | Non-PD |
| Female | 81 | White | 6 | Non-PD |
| Male | 88 | White | 8 | Non-PD |
| Male | 79 | White | 8.72 | Non-PD |
| Male | 86 | White | 7.87 | Non-PD |
| Male | 77 | Black or African | 3.47 | Non-PD |
| Male | 79 | White | 5.92 | Non-PD |
| Male | 76 | White | 2.92 | Non-PD |
| Male | 78 | White | 8.08 | Non-PD |
| Male | 84 | White | 2.92 | Non-PD |
| Male | 75 | Black or African | 78.17 | Non-PD |
| Female | 87 | Unknown | 5.58 | PD |
| Female | 77 | White | 11.25 | PD |
| Female | 73 | Unknown | 19.25 | PD |
| Female | 85 | Unknown | 13.58 | PD |
| Female | 79 | Unknown | 21.12 | PD |
| Female | 78 | White | 7 | PD |
| Female | 82 | White | 8 | PD |
| Female | 78 | White | 9.67 | PD |
| Female | 76 | White | 3.83 | PD |
| Male | 88 | White | 5.25 | PD |
| Male | 75 | Unknown | 5.33 | PD |
| Male | 84 | White | 4 | PD |
| Male | 77 | White | 3.98 | PD |
| Male | 77 | White | 8.42 | PD |
| Male | 81 | White | 4.5 | PD |
| Male | 77 | White | 2.27 | PD |

**Table 2**. Mitochondrial dynamics-related proteins and RNA adjusted for age and sex.

| **Proteins** | **PD/Non-PD (Adjusted p-value)** | **Age (p-value)** | **Sex/Male (p-value)** |
| --- | --- | --- | --- |
| DRP1 | 0.040 * | 0.80 | 0.40 |
| OPA-1 | 0.040 * | 0.20 | 0.12 |
| MiD51 | 0.047 * | 0.70 | 0.80 |
| MFN2 | 0.300 | 0.20 | 0.90 |
| **Genes** | **PD/Non-PD (Adjusted p-value)** | **Age (p-value)** | **Sex/Male (p-value)** |
| *DNM1L* | 0.001 * | 0.20 | 0.30 |
| *OPA-1* | 0.001 * | 0.20 | 0.60 |
| *MFF* | 0.051 * | 0.20 | 0.80 |
| *MIEF1* | 0.025 * | 0.30 | 0.06 |
| *MIEF2* | 0.023 * | 0.60 | 0.14 |
| *MFN2* | 0.088 | 0.60 | 0.60 |
| * Significant p-values after sex and age adjustment | | | |

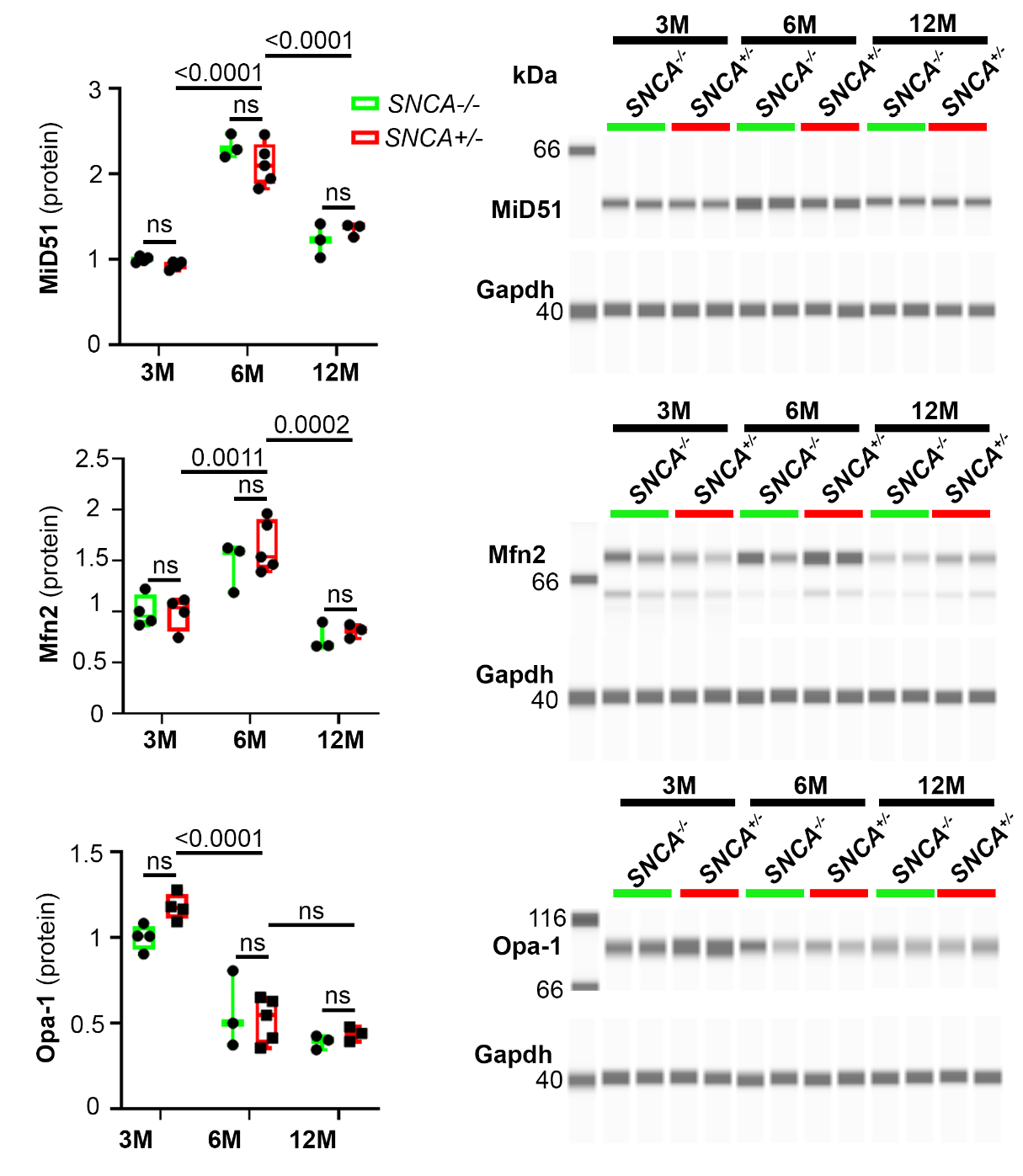

**Fig. S1** Age dependent changes in mitochondrial dynamics. Immunoblotting for MiD51, Mfn2, and Opa-1 in ventral midbrains of 3-, 6-, and 12-month-old *SNCA* mice and their WT littermates. Data represents ± SEM, normalized to Gapdh. Age-dependent and genotype-dependent changes were analyzed using two-way ANOVA followed by Tukey’s post hoc tests. Genotypic changes independent of age were analyzed using unpaired t-test, n=3-5.

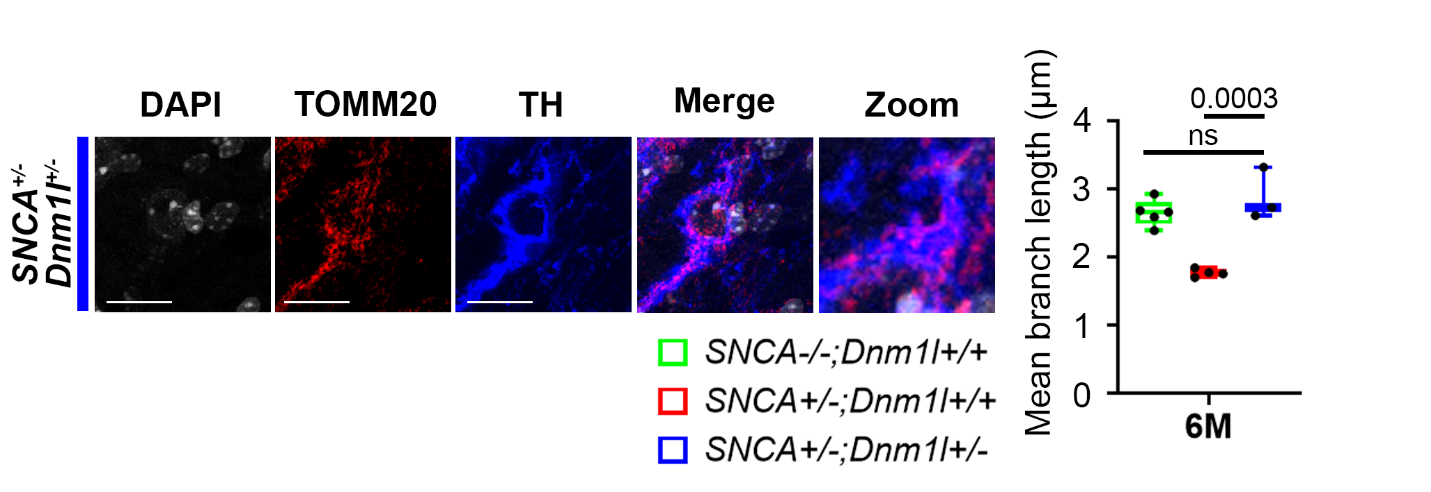

**Fig. S2** Immunofluorescent staining of TOMM20 to visualize mitochondrial morphology in 6-month-old *SNCA* mice with heterozygous Drp1-KO and their WT littermate counterparts. Data represent mean ± SEM, One-way ANOVA, n=3-5

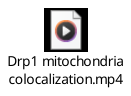

**Fig. S3** IMARIS 3D rendering video conceptualizing Drp1 mitochondria colocalization analysis
